# SARS-CoV-2 neutralizing serum antibodies in cats: a serological investigation

**DOI:** 10.1101/2020.04.01.021196

**Authors:** Qiang Zhang, Huajun Zhang, Kun Huang, Yong Yang, Xianfeng Hui, Jindong Gao, Xinglin He, Chengfei Li, Wenxiao Gong, Yufei Zhang, Cheng Peng, Xiaoxiao Gao, Huanchun Chen, Zhong Zou, Zhengli Shi, Meilin Jin

**Author notes:** Corresponding author. E-mail address (Meilin Jin); (Zhong Zou); (Zhengli Shi). These authors have contributed equally to this work (Qiang Zhang, Huajun Zhang).

## Abstract

Coronavirus disease 2019 (COVID-19) caused by severe acute respiratory syndrome coronavirus 2 (SARS-CoV-2) was first reported in Wuhan, China, and rapidly spread worldwide. Previous studies suggested cat could be a potential susceptible animal of SARS-CoV-2. Here, we investigated the infection of SARS-CoV-2 in cats by detecting specific serum antibodies. A cohort of serum samples were collected from cats in Wuhan, including 102 sampled after COVID-19 outbreak, and 39 prior to the outbreak. 15 of 102 (14.7%) cat sera collected after the outbreak were positive for the receptor binding domain (RBD) of SARS-CoV-2 by indirect enzyme linked immunosorbent assay (ELISA). Among the positive samples, 11 had SARS-CoV-2 neutralizing antibodies with a titer ranging from 1/20 to 1/1080. No serological cross-reactivity was detected between the SARS-CoV-2 and type I or II feline infectious peritonitis virus (FIPV). Our data demonstrates that SARS-CoV-2 has infected cat population in Wuhan during the outbreak.

## Introduction

In December, 2019, an outbreak of pneumonia of unknown cause occurred in Wuhan, China. The pathogen was soon identified to be the severe acute respiratory syndrome coronavirus 2 (SARS-CoV-2), and the disease was designated coronavirus disease 2019 (COVID-19) by World Health Organization (WHO) (Chen et al., 2020; Zhou et al., 2020a). The clinical symptoms of COVID-19 mainly include asymptomatic infection, mild-to-severe respiratory tract illness, and even death (Huang et al., 2020). Compared with SARS-CoV, SARS-CoV-2 has the higher basic reproduction number, representing more transmissibility (Liu et al., 2020a). Within a very short period of time, COVID-19 has quickly become a very serious threat to travel, commerce, and human health in the worldwide (Bernard Stoecklin et al., 2020; Ghinai et al., 2020; Iacobucci, 2020; Shim et al., 2020; Tuite et al., 2020). By Mar. 28, 2020, a total of 512,701 confirmed cases, including 23,495 deaths (4.58%), involving 202 countries, areas or territories, have been reported globally by WHO (https://www.who.int/emergencies/diseases/novel-coronavirus-2019).

The outbreak of COVID-19 was first confirmed in Wuhan, China, possibly associated with a seafood market. However, so far, there is no evidence that the seafood market is the original source of SARS-CoV-2 (Guo et al., 2020). Before SARS-CoV-2, 4 types of beta coronaviruses can infect humans, including SARS-CoV and MERS-CoV which are highly pathogenic and both originated from bats (Li et al., 2020a; Liu et al., 2020b). Genome analysis showed that SARS-CoV-2 has 96.2% overall genome sequence identity with Bat CoV RaTG13, indicating that SARS-CoV-2 could also originate from bats (Zhou et al., 2020b). The transmission of SARS-CoV-2 from bats to humans was suspected to via the direct contact between humans and intermediate host animals (Guo et al., 2020). However, it remains unclear which animals were the intermediate host of SARS-CoV-2. Our previous study showed that SARS-CoV-2 uses the same cell entry receptor, angiotensin converting enzyme II (ACE2), as SARS-CoV (Zhou et al., 2020b), suggesting that SARS-CoV-2 has the same host range as SARS-CoV. Previous report demonstrated that SARS-CoV can infect ferrets and cats (Martina et al., 2003), implying that they might be also susceptible to SARS-CoV-2.

As one of the most popular pets, cats have very close contact with humans. Therefore, it is very important to investigate the prevalence of SARS-CoV-2 in cat, especially in the outbreak areas. However, there is no survey about the prevalence of SARS-CoV-2 in cats so far. Serological studies are suitable for the screening of antibody against the SARS-CoV-2 in animals (Reusken et al., 2013). At present, several methods have been applied for the antibody test of SARS-CoV-2 in human (Li et al., 2020b). However, there is no available method for the detection of cat antibody against the SARS-CoV-2. Here, we investigate the serological prevalence of SARS-CoV-2 in cats by an indirect ELISA and virus neutralization test, providing the first evidence of SARS-CoV-2 infection in cats.

## Results

A total of 143 cat sera were screened by indirect enzyme linked immunosorbent assay (ELISA) for antibody reactivity against recombinant receptor binding domain (RBD) of SARS-CoV-2 spike protein. From the 39 sera collected before the outbreak, whose OD varied from 0.091 to 0.261, we set the cut-off as 0.32. As shown in Table 1 and Figure 1, 15 cat sera (14.7%) collected after the outbreak were positive, with five strong positive ones with OD more than 0.6. Both type I and II feline infectious peritonitis virus (FIPV) hyperimmune sera showed no cross-reactivity with SARS-CoV-2 RBD protein.

**Table 1.** COVID-19 patient contact histories of the ELISA positive cats.

| **Cat** | **ELISA ( OD450 )** | **Neutralization titer** | **Background** |
| --- | --- | --- | --- |
| #1 | 0.353 | 1/40 | from pet hospital |
| #2 | 0.334 | 1/80 | stray cat |
| #3 | 0.348 | 1/40 | stray cat |
| #4 | 0.687 | 1/360 | owner COVID-19 patient |
| #5 | 0.394 | 1/40 | from pet hospital |
| #6 | 0.401 | — | from pet hospital |
| #7 | 0.379 | — | from pet hospital |
| #8 | 0.345 | 1/20 | from pet hospital |
| #9 | 0.351 | 1/40 | from pet hospital |
| #10 | 0.624 | 1/20 | stray cat |
| #11 | 0.342 | 1/40 | stray cat |
| #12 | 0.852 | — | stray cat |
| #13 | 0.437 | — | stray cat |
| #14 | 1.432 | 1/360 | owner COVID-19 patient |
| #15 | 1.095 | 1/1080 | owner COVID-19 patient |

**Figure 1.**
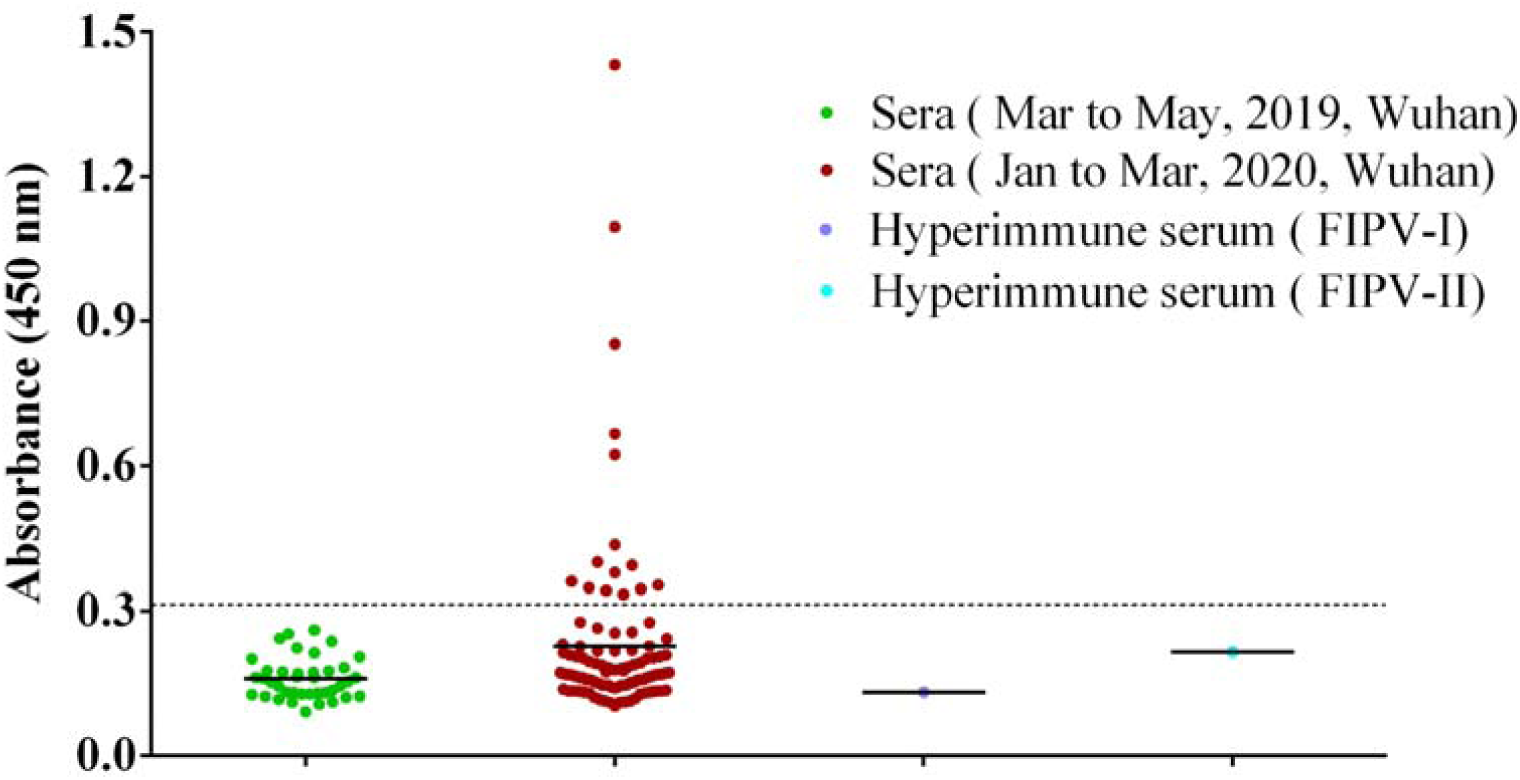
ELISA of cat serum samples against the recombinant receptor binding domain (RBD) of SARS-CoV-2 spike. The dashed line is the positive cut-off. Each dot represents one individual sample within each antigen panel.

To confirm the presence of SARS-CoV-2 specific antibody in cat sera, all of 15 ELISA positive sera were tested by virus neutralization tests (VNT) for SARS-CoV-2. Of which, 11 cat sera had SARS-CoV-2 neutralizing antibodies with a titer ranging from 1/20 to 1/1080 (Table 1, Figure 2A). However, 4 sera including #12, which was ELISA strong positive with OD of 0.85, showed no neutralizing activity. And another ELISA strong positive, #10, had very weak neutralizing activity. But strong neutralization was observed for the other three ELISA strong positive sera, #4, #14 and #15, with neutralizing titer of 1/360 to 1/1080. Consistent with the high neutralizing titer, the owners of Cat#4, Cat#14 and Cat#15 were diagnosed as COVID-19 patients. Cat#1, Cat#5∼9 was from pet hospitals, while Cat#2, Cat#10∼13 were initially stray cats and kept in animal protection shelters after the outbreak. Again, both type I and II FIPV hyperimmune sera were negative for VNT.

**Figure 2.**
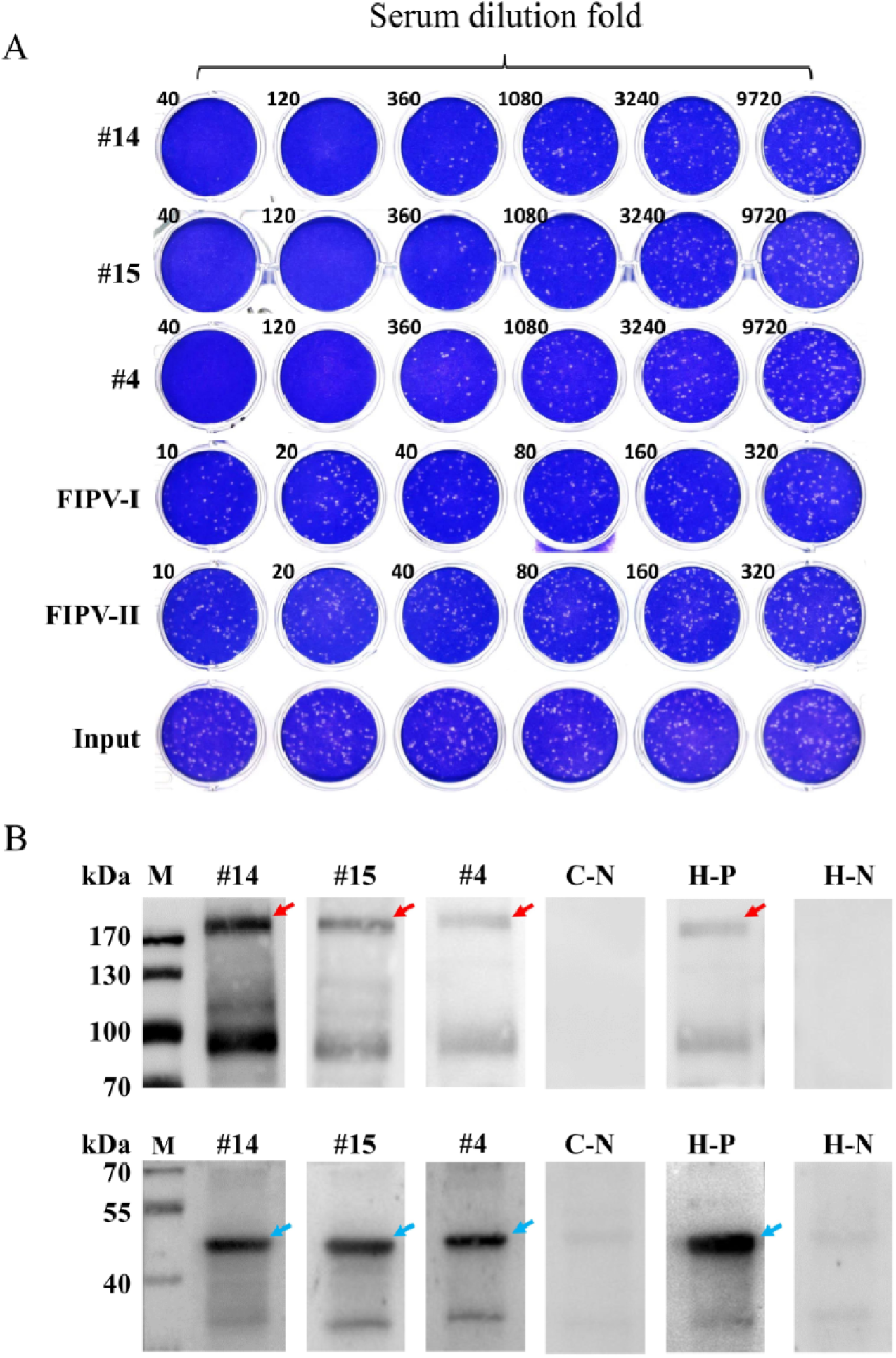
Virus neutralization test and western blotting assay of cat serum samples for SARS-CoV-2. **(A)** Morphology of SARS-CoV-2 viral plaques. Three representative sera are shown (#4, #14 and #15 corresponding to cat ID numbers in table 1) as well as hyperimmune sera of type I and II FIPV, and the virus input control. **(B)** Western blotting assay of cat or human serum samples for SARS-CoV-2. The convalescent serum of COVID-19 patient was used as a positive control. The negative cat serum of ELISA or healthy human serum was used as negative control. All of the detected serum samples were used at a dilution of 1:100. C-N, negative cat serum. H-P, human convalescent serum. H-N, healthy human serum. Red arrows, S protein. Blue arrows, N protein.

Western blot assay was also performed to further verify the existence of SARS-CoV-2 specific IgG in cat serum. As shown in Figure 2 B, #4, #14 and #15 sera detected S and N proteins of purified SARS-CoV-2, as human convalescent serum. In contrast, the ELISA negative cat serum and healthy human serum didn’t probe the protein bands.

## Discussion

In this study, we detected the presence of SARS-CoV-2 antibodies in cats in Wuhan during the COVID-19 outbreak with ELISA, VNT and western blot. A total of 102 cats were tested, 15 (14.7%) were positive for RBD based ELISA and 11 (10.8%) were further positive with VNT. These results demonstrated that SARS-CoV-2 has infected cat populations in Wuhan, implying that this risk could also occur at other outbreak regions. Retrospective investigation confirmed that all of ELISA positive sera were sampled after the outbreak, suggesting that the infection of cats could be due to the virus transmission from humans to cats. Certainly, it is still needed to be verified via investigating the SARS-CoV-2 infections before this outbreak in a wide range of sampling. At present, there is no evidence of SARS-CoV-2 transmission from cats to humans. However, a latest report shows that SARS-CoV-2 can transmit between cats via respiratory droplets (Hualan Chen, 2020), so, a strong warning and regulations still should be issued to block this potential transmission route.

The three cats owned by COVID-19 patients had the highest neutralization titer (1/360, 1/360, and 1/1080, respectively). On the contrary, the sera collected from pet hospital cats and stray cats had neutralizing activity of 1/20 to 1/80, indicating that the high neutralization titers could be due to the close contact between cats and COVID-19 patients. Although the infection in stray cats was not fully understood, it is reasonable to speculate that these infections are probably due to the contact with SARS-CoV-2 polluted environment, or COVID-19 patients who fed the cats.

In addition, we also collected nasopharyngeal and anal swabs of each cat, and conducted SARS-CoV-2 specific qRT-PCR using a commercial kit which targeted *ORF1ab* and *N* genes. However, no double gene positive sample was detected. The reason might be (1) that the viral RNA load is too low to be detected; (2) as SARS-CoV (Martina et al., 2003), the period that cat shed SARS-CoV-2 may be very short, along with asymptomatic infection, we didn’t catch the moment of acute infection; (3) there may be variants in the genomic sequences in cats, leading to the failure in amplification in cat samples.

To the best of our knowledge, this is the first report that animals produce specific neutralizing antibodies against SARS-CoV-2 under natural conditions. Our study pointed out the risk of cats involved in the transmission of SARS-CoV-2. More studies are needed to investigate the transmission route of SARS-CoV-2 from humans to cats. Importantly, an immediate action should be implemented to keep in a suitable distance between humans and companion animals such as cats and dogs, and strict hygiene and quarantine measures should also be carried out for these animals.

## Acknowledgments

We acknowledge Jiangxia Tongji hospital for providing the convalescent serum of COVID-19 patient. We thank Professor Guiqing Peng (Huazhong Agriculture University) for providing the hyperimmune sera against type I and II FIPV.

## Author Contributions

M.J., Z.Z., Z.S., Q.Z., and H.Z. conceived and designed the study, wrote the report, drew the figures, and generated, analyzed, and interpreted data. J.G., X.H., and C.L. collected the samples. Q.Z., H.Z., K.H., Y.Y., X.H., W.G., Y.Z., C.P., and X.G. performed the experiments. H.C. coordinated the study. All authors critically revised the manuscript for important intellectual content and gave final approval for the version to be published. All authors agree to be accountable for all aspects of the work in ensuring that questions related to the accuracy or integrity of any part of the work are appropriately investigated and resolved.

## Declaration of interests

The authors declare no competing interests.

## Methods

### Sample collection

A total of 102 cats were sampled from animal shelters or pet hospitals of Wuhan between Jan. and Mar., 2020. Blood samples were collected via leg venipuncture and sera were separated and stored at −20°C until further processing. Nasopharyngeal and anal swabs were collected and put into tubes containing viral transport medium-VTM (Copan Diagnostics, Brescia, Italy) (Haagmans et al., 2014). All samples were collected under full personal-protective equipment, including head covers, goggles, N95 masks, gloves, and disposable gowns. A set of 39 cat sera were retrieved from the serum bank in our lab, which were collected from Wuhan between Mar. and May, 2019. Hyperimmune sera were obtained from Neuropathy Pathogen Laboratory, Huazhong Agriculture University, with neutralization titers of 1/640 and 1/1280, respectively, against type I and II feline infectious peritonitis virus (FIPV). The convalescent serum of a COVID-19 patient was collected from Jiangxia Tongji hospital with the consent of the patient and a neutralization titer 1/1280.

### Virus and cells

SARS-CoV-2 (IVCAS 6.7512) was isolated from a COVID-19 patient as previously described.(Zhou et al., 2020b) Vero E6 was purchased from ATCC (ATCC® CRL-1586™).

### Enzyme-linked immunosorbent assay (ELISA)

Antibody was tested by indirect ELISA with the SARS-CoV-2 RBD protein (Sino Biological Inc., China) and peroxidase conjugated goat anti-cat IgG (Sigma-Aldrich, USA). Briefly, ELISA plates were coated overnight at 4 □ with RBD protein (1 μg/ml, 100μl per well). After blocked with PBS containing 5% skim milk for 2 h at 37 □, the plates were added with sera at a dilution of 1: 40. After incubation for 30 min at 37 °C, the plates were washed 5 times with washing buffer (PBS containing 0.05% Tween-20). A 1:20000 diluted anti-cat IgG was added and incubated for an additional 30 min. After another 5 washes, TMB Substrate (Sigma-Aldrich, USA) was added and incubated for 10 min. Then the reaction was stopped, and optical density (OD) was measured at 450 nm. Those sera were considered positive if the OD values were twice higher than the mean OD of the 39 sera collected between Mar and May, 2019.

### Virus neutralization test (VNT)

For virus neutralization test, serum samples were heat-inactivated by incubation at 56°C for 30 min. Each serum sample was serially diluted with Dulbecco’s Modified Eagle Medium (DMEM) as two fold or three fold according to the OD value and the sample quality, mixed with equal volume of diluted virus and incubated at 37°C for 1 h. Vero E6 cells in 24-well plates were inoculated with the sera-virus mixture at 37°C; 1 h later, the mixture was replaced with DMEM containing 2.5% FBS and 0.8% carboxymethylcellulose. The plates were fixed with 8% paraformaldehyde and stained with 0.5% crystal violet 3 days later. All samples were tested in duplicate and neutralization titers were defined as the serum dilution resulting in a plaque reduction of at least 50% (Davies et al., 2005).

### Western blotting assay

The total protein concentration of purified and inactivated SARS-CoV-2 was determined by Bradford protein assay (Su et al., 2018). 4 μg protein was subjected to 8% sodium dodecyl sulfate-polyacrylamide gel electrophoresis (SDS-PAGE) and transferred on to nitrocellulose membrane. Then viral proteins were blotted with cat sera or human convalescent serum. Protein bands were visualized by incubation with a goat anti-cat IgG or mouse anti-human IgG and then detected using the ECL System (Amersham Life Science, Arlington Heights, IL, USA).

## References

Bernard Stoecklin, S., Rolland, P., Silue, Y., Mailles, A., Campese, C., Simondon, A., Mechain, M., Meurice, L., Nguyen, M., Bassi, C., et al. (2020). First cases of coronavirus disease 2019 (COVID-19) in France: surveillance, investigations and control measures, January 2020. Euro surveillance : bulletin Europeen sur les maladies transmissibles = European communicable disease bulletin 25.

Chen, N., Zhou, M., Dong, X., Qu, J., Gong, F., Han, Y., Qiu, Y., Wang, J., Liu, Y., Wei, Y., et al. (2020). Epidemiological and clinical characteristics of 99 cases of 2019 novel coronavirus pneumonia in Wuhan, China: a descriptive study. Lancet 395, 507–513.

Davies, D.H., McCausland, M.M., Valdez, C., Huynh, D., Hernandez, J.E., Mu, Y., Hirst, S., Villarreal, L., Felgner, P.L., and Crotty, S. (2005). Vaccinia virus H3L envelope protein is a major target of neutralizing antibodies in humans and elicits protection against lethal challenge in mice. Journal of virology 79, 11724–11733.

Ghinai, I., McPherson, T.D., Hunter, J.C., Kirking, H.L., Christiansen, D., Joshi, K., Rubin, R., Morales-Estrada, S., Black, S.R., Pacilli, M., et al. (2020). First known person-to-person transmission of severe acute respiratory syndrome coronavirus 2 (SARS-CoV-2) in the USA. Lancet.

Guo, Y.R., Cao, Q.D., Hong, Z.S., Tan, Y.Y., Chen, S.D., Jin, H.J., Tan, K.S., Wang, D.Y., and Yan, Y. (2020). The origin, transmission and clinical therapies on coronavirus disease 2019 (COVID-19) outbreak - an update on the status. Military Medical Research 7, 11.

Haagmans, B.L., Al Dhahiry, S.H., Reusken, C.B., Raj, V.S., Galiano, M., Myers, R., Godeke, G.J., Jonges, M., Farag, E., Diab, A., et al. (2014). Middle East respiratory syndrome coronavirus in dromedary camels: an outbreak investigation. The Lancet infectious diseases 14, 140–145.

Hualan Chen. Susceptibility of ferrets, cats, dogs, and different domestic animals to SARS-coronavirus-2. doi: https://doi.org/10.1101/2020.03.30.015347.

Huang, C., Wang, Y., Li, X., Ren, L., Zhao, J., Hu, Y., Zhang, L., Fan, G., Xu, J., Gu, X., et al. (2020). Clinical features of patients infected with 2019 novel coronavirus in Wuhan, China. Lancet 395, 497–506.

Iacobucci, G. (2020). Covid-19: all non-urgent elective surgery is suspended for at least three months in England. Bmj 368, m1106.

Li, B., Si, H.R., Zhu, Y., Yang, X.L., Anderson, D.E., Shi, Z.L., Wang, L.F., and Zhou, P. (2020a). Discovery of Bat Coronaviruses through Surveillance and Probe Capture-Based Next-Generation Sequencing. mSphere 5.

Li, Z., Yi, Y., Luo, X., Xiong, N., Liu, Y., Li, S., Sun, R., Wang, Y., Hu, B., Chen, W., et al. (2020b). Development and Clinical Application of A Rapid IgM-IgG Combined Antibody Test for SARS-CoV-2 Infection Diagnosis. Journal of medical virology.

Liu, Y., Gayle, A.A., Wilder-Smith, A., and Rocklov, J. (2020a). The reproductive number of COVID-19 is higher compared to SARS coronavirus. Journal of travel medicine 27.

Liu, Z., Xiao, X., Wei, X., Li, J., Yang, J., Tan, H., Zhu, J., Zhang, Q., Wu, J., and Liu, L. (2020b). Composition and divergence of coronavirus spike proteins and host ACE2 receptors predict potential intermediate hosts of SARS-CoV-2. Journal of medical virology.

Martina, B.E., Haagmans, B.L., Kuiken, T., Fouchier, R.A., Rimmelzwaan, G.F., Van Amerongen, G., Peiris, J.S., Lim, W., and Osterhaus, A.D. (2003). Virology: SARS virus infection of cats and ferrets. Nature 425, 915.

Reusken, C.B., Haagmans, B.L., Muller, M.A., Gutierrez, C., Godeke, G.J., Meyer, B., Muth, D., Raj, V.S., Smits-De Vries, L., Corman, V.M., et al. (2013). Middle East respiratory syndrome coronavirus neutralising serum antibodies in dromedary camels: a comparative serological study. The Lancet infectious diseases 13, 859–866.

Shim, E., Tariq, A., Choi, W., Lee, Y., and Chowell, G. (2020). Transmission potential and severity of COVID-19 in South Korea. International journal of infectious diseases : IJID : official publication of the International Society for Infectious Diseases.

Su, X., Tian, Y., Zhou, H., Li, Y., Zhang, Z., Jiang, B., Yang, B., Zhang, J., and Fang, J. (2018). Inactivation Efficacy of Nonthermal Plasma-Activated Solutions against Newcastle Disease Virus. Applied and environmental microbiology 84.

Tuite, A.R., Ng, V., Rees, E., and Fisman, D. (2020). Estimation of COVID-19 outbreak size in Italy. The Lancet infectious diseases.

Zhou, F., Yu, T., Du, R., Fan, G., Liu, Y., Liu, Z., Xiang, J., Wang, Y., Song, B., Gu, X., et al. (2020a). Clinical course and risk factors for mortality of adult inpatients with COVID-19 in Wuhan, China: a retrospective cohort study. Lancet.

Zhou, P., Yang, X.L., Wang, X.G., Hu, B., Zhang, L., Zhang, W., Si, H.R., Zhu, Y., Li, B., Huang, C.L., et al. (2020b). A pneumonia outbreak associated with a new coronavirus of probable bat origin. Nature 579, 270–273.

